# Quantitative phase deformability cytometry (QP-DC) for precise and clinically relevant multiparametric immune cell profiling

**DOI:** 10.1101/2025.10.27.684875

**Authors:** Kyoohyun Kim, Eoghan N. O’Connell, Christine Schauer, Janina Schoen, Jiwoo Shim, Florian Mayerle, Philipp Radler, Philipp Lebhardt, Martin Kräter, Jürgen Rech, Jens Langejürgen, Martin Herrmann, Jochen Guck

**Author notes:** M.H. and J.G. equally contributed to senior authorship.

## Abstract

Imaging flow cytometry enables the detailed analysis of cell morphology and internal structures through high-throughput cell imaging, and quantitative phase imaging (QPI)-based microfluidic approaches have extended this by providing label-free measures such as dry mass and refractive index (RI). Building on these developments, we present quantitative phase deformability cytometry (QP-DC), which integrates QPI with deformability cytometry to simultaneously measure morphology, mechanics, and intrinsic biophysical parameters such as mass density and dry mass. Numerical refocusing ensures in-focus images independent of axial position, improving precision in contour detection and feature extraction. Using microspheres and whole blood, we validated QP-DC and then applied it to neutrophils under lipopolysaccharide (LPS) stimulation and from patients with systemic lupus erythematosus (SLE). QP-DC revealed LPS-induced reductions in neutrophil mass density and identified heterogeneous subpopulations in SLE. These results demonstrate the capability of QP-DC for precise biophysical and mechanical characterization, offering significant potential for research and clinical diagnostics.

## Introduction

Imaging flow cytometry has been developed for measuring biological and physical properties of a large number of cells. It employs high-throughput acquisition of single-cell images of samples flowing through microfluidic channels^1–3^. It provides image-based analysis of spatially-resolved characteristics such as cell morphology and internal structure distributions. It thus enables rapid diagnoses of disease conditions and high-resolution drug screening. Recently, real-time deformability cytometry (RT-DC), a specific implementation of imaging flow cytometry, has explored rapid and high-throughput assessment of cell deformability in addition to cell morphology^4^, which provides critical insights into cell mechanics under various physiological and pathological states^5–7^.

One of the key applications of imaging flow cytometry is the analysis of blood cells, particularly neutrophils, which are central to the innate immune response. Recent studies have revealed substantial heterogeneity within neutrophil populations under different physiological and pathological conditions, including the emergence of low-density neutrophils (LDNs)—a subpopulation characterized by reduced buoyant density and altered functional reactivity. LDNs were initially described in patients with systemic lupus erythematosus (SLE), in which they comprise a substantial part of the circulating neutrophils and contribute to disease pathology through pro-inflammatory functions and spontaneous intravascular neutrophil extracellular trap (NET) formation^8–10^. Beyond SLE, LDNs have been associated with various disease states, yet their precise biological origin and functional significance remain elusive^8,11^. Imaging flow cytometry and RT-DC have recently enabled high-throughput analysis of neutrophil morphology and mechanics, providing new insight into LDN phenotypes^12,13^.

To perform image-based analysis, conventional imaging flow cytometry involves acquiring bright field and fluorescence images, followed by image processing to extract key features such as cell contour, area, and brightness. The accuracy and precision of these processes are highly dependent on the expertise of operators maintaining optimal experimental conditions, including proper optical focus and adequate brightness contrast between the sample and background^1,14^. Achieving consistent focus is still challenging, as it is susceptible to variations in sample size, mechanical vibrations, thermal expansion of the microfluidic chip, and flow instabilities. Moreover, the inherent axial distribution of objects within the channel means that a fixed optical focus often results in some objects being out of focus. These factors introduce variability and potential errors in image acquisition and image-based analysis in imaging flow cytometry, highlighting the need for more reliable methods to maintain optimal experimental conditions.

In this study, we introduce quantitative phase deformability cytometry (QP-DC), which integrates quantitative phase imaging (QPI) with deformability cytometry to address the limitations of conventional imaging flow cytometry. QPI utilizes interferometric techniques to measure complex optical fields consisting of both the amplitude and phase map^15,16^. We employed self-reference interferometry to acquire complex optical fields from samples flowing through microfluidic channels using a high-speed camera. Numerical focusing based on Rayleigh-Sommerfeld propagation can generate in-focus images of the samples during post-processing. This numerical focusing capability significantly improves the precision of extracting cell morphology and deformability independent of axial positions. Furthermore, the phase maps measured by QPI provide additional biophysical parameters including dry mass, refractive index (RI), and mass density. These parameters are increasingly recognized for their roles in biological processes such as differentiation, cell growth, protein synthesis, and condensate formation^17^.

We validated the QP-DC method using a mixture of microspheres with the same diameter but different RI values, and characterized the mechanical and biophysical properties of different blood cell types from whole blood. To demonstrate the clinical utility of QP-DC, we examined neutrophil biophysical changes during activation and disease progression. Upon stimulation with lipopolysaccharide (LPS), neutrophils from healthy donors showed increased deformability and reduced mass density, primarily driven by volume expansion. These effects were almost completely suppressed by phloretin, confirming the role of aquaporin-mediated water influx. QP-DC measurements from SLE patients revealed reduced neutrophil mass density and increased area, consistent with the presence of LDNs previously linked to disease severity. QP-DC measurements revealed a trend between decreasing mass density and SLE Disease Activity Index (SLE-DAI) scores, alongside morphological signs of early NET formation. These findings establish QP-DC as a rapid, label-free method for quantifying disease-associated neutrophil phenotypes.

In summary, QP-DC enables high-throughput, quantitative analysis of cell mechanics and intrinsic physical properties, overcoming key limitations of conventional imaging flow cytometry. By combining precise morphological and biophysical profiling, it offers a powerful method for studying immune cell heterogeneity in both research and clinical contexts.

## Results

### Quantitative phase deformability cytometry (QP-DC) setup

The experimental setup for QP-DC utilized self-reference interferometry to image samples flowing within a microfluidic channel with the channel width of 20 μm (Fig. 1a, see also Materials and Methods). When samples flow through a microfluidic channel with a channel width slightly larger than sample size, the samples are deformed in a contactless manner by hydrodynamic shear and normal stresses^4,18^. A collimated laser beam (*λ* = 520 nm) illuminated samples flowing through the microfluidic channel, and the sample images were collected by an objective lens and a tube lens. A Rochon polarizer in the self-reference interferometry duplicated the sample image into two beams (ordinary and extraordinary). The lateral shearing of these beams at the camera plane generated an off-axis hologram, which was recorded with a high-speed camera with the frame rate of 3,000 Hz (Fig. 1b). We chose a region of interest of the hologram generated by the interference between the sample image in the ordinary beam and the background outside the microfluidic channel in the extraordinary beam. Conventional self-reference interferometry techniques have required sparse samples distribution to avoid image overlapping between the ordinary and extraordinary beams^19–21^, but QP-DC overcame this constraint due to the spatial confinement by the microfluidic channel. Consequently, the lateral shearing distance between duplicated beams was minimized to the channel width (20 μm) and did not require sparse sample distribution. From the measured off-axis holograms, the complex optical fields consisting of amplitude and phase were retrieved by a field retrieval algorithm based on Fourier transform^22^, and clearly showed samples flowing inside the microfluidic channel (Fig. 1c). The individual samples in the retrieved phase maps were segmented by thresholding based on phase values, and the morphological and mechanical parameters including cell area and contour length were extracted by convex hull calculations for each segmentation, which is described elsewhere^14^. Additionally, the dry mass, RI, and mass density were quantitatively characterized from the phase maps (see Materials and Methods).

**Figure 1.**
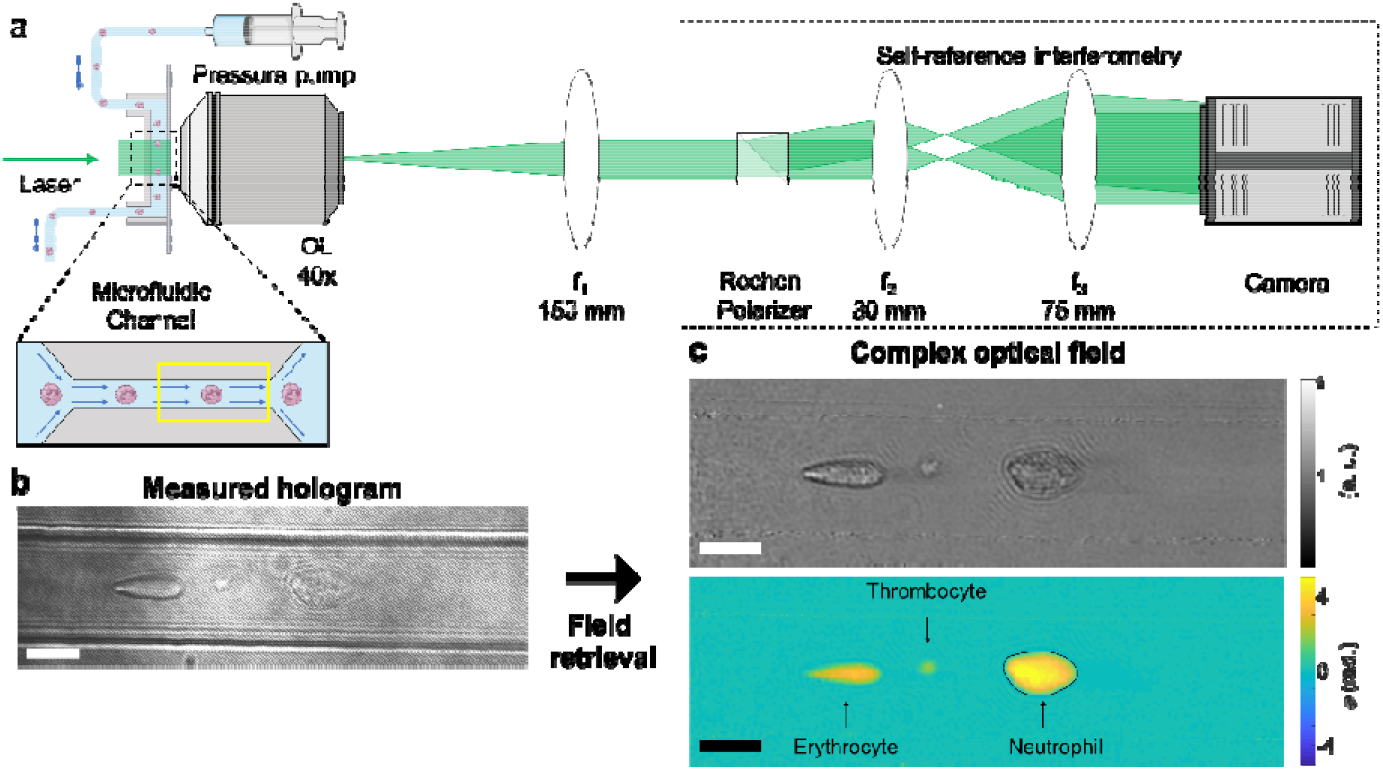
Quantitative phase deformability cytometry (QP-DC). (**a**) Schematic of the experimental setup. (**b**) A measured hologram with examples of different blood cells. (**c**) Complex optical field calculated from (**b**), consisting of amplitude (top) and phase (bottom) maps, with blood cell types indicated. The contour of the example neutrophil is highlighted in a dashed line, which is used to extract morphological and physical parameters of each samples including deformation, area, dry mass, and mass density. Scale bars: 10 μm. Fig. 1**a** was created with BioRender.

### Phase-based numerical refocusing enhances data precision

QP-DC enhances precision of imaging flow cytometry by providing numerical refocusing of acquired holograms, which is enabled by its capability to measure both the amplitude and phase of light through QPI. This aspect is especially crucial for accurate extraction of sample contours from images that may experience axial focus drift due to various factors, including sample size variations, mechanical vibrations, thermal expansion of the microfluidic chip, and flow instabilities. This focal drift can affect the accuracy of drawing sample contours and consequently image-based feature extraction. QP-DC mitigated this issue through numerical refocusing. Using Rayleigh-Sommerfeld backpropagation of the acquired complex optical field^23,24^, we generated an axial stack of propagated complex optical fields ranging from z = −10 μm to 10 μm with a step size of 0.2 μm. The optimal optical focus was determined using a defined focus criterion (See Materials and Methods and Supplementary Figure 1).

We validated the numerical focusing capabilities of QP-DC using a mixture of poly(methyl methacrylate) (PMMA) and silicon dioxide (silica) microspheres, each 4 μm in diameter but with different RI values (*n*_PMMA_ = 1.4945, *n*_Silica_ = 1.4613), suspended in a 40% (w/w) sucrose solution. Although the microspheres experience laminar flow in the microfluidic channel, they can lie at different axial planes. For example, in Fig. 2a, silica and PMMA microspheres are found at different focuses, even within the same field of view. The numerical refocusing correctly determined the axial positions as z = −2.4 μm for the silica and 1.4 μm for PMMA at the frame (Fig. 2b and Supplementary Figure 1). The refocused amplitude and phase maps show that the complex optical fields of each microsphere were accurately refocused, with minimized diffraction patterns at the sample boundaries.

**Figure 2.**
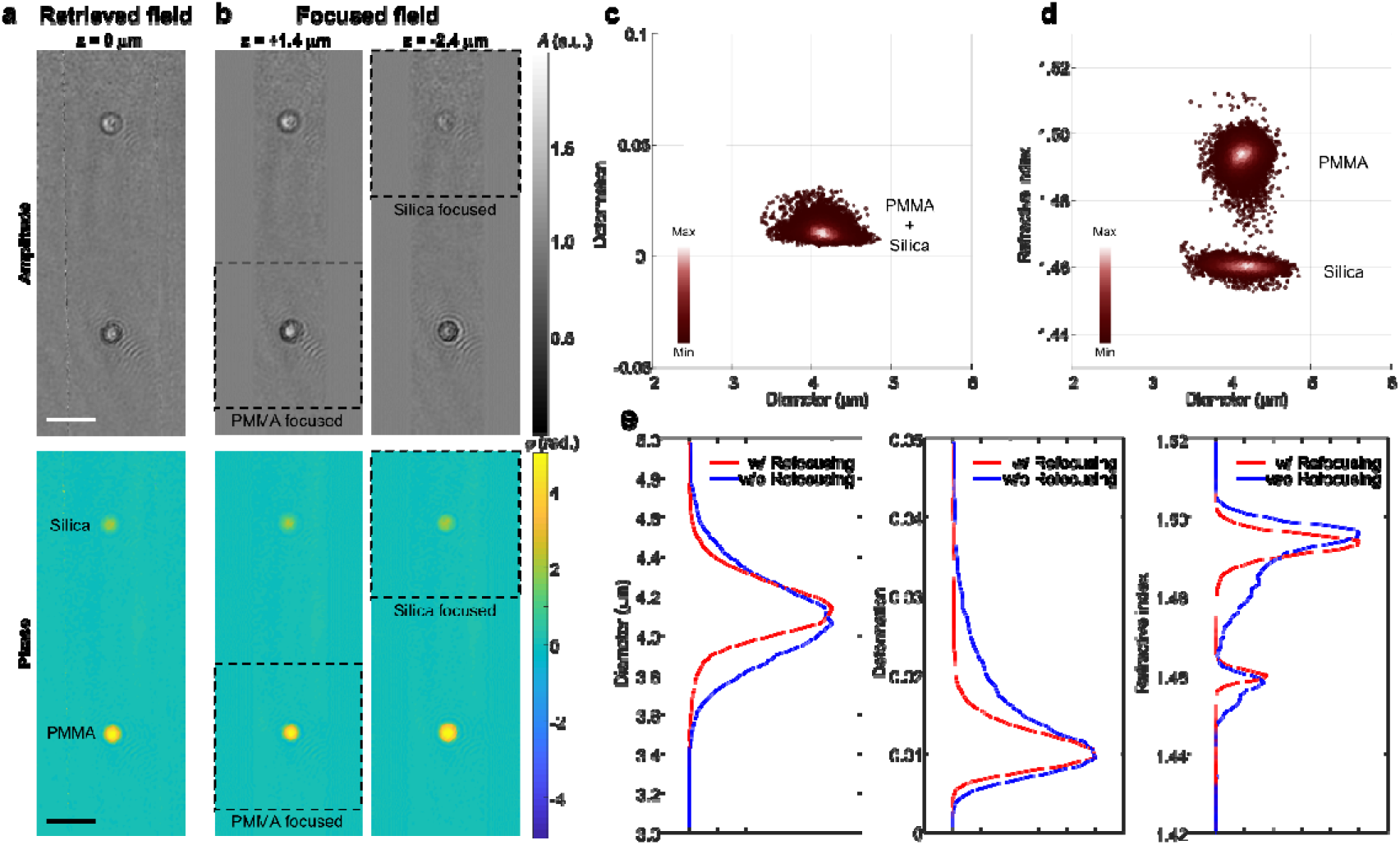
Numerical refocusing. (**a**) The amplitude (top) and phase (bottom) of a silica and PMMA bead retrieved from the acquired hologram. (**b**) The amplitude (top) and phase (bottom) maps of a PMMA (left) and a silica (right) microsphere at the focused plane found at z = 1.4 μm and −2.4 μm, respectively. The dashed boxes indicate the focused microspheres. (**c** – **d**) 2D scatter plots of (**c**) deformation and (**d**) RI versus diameter for a mixture of silica and PMMA microspheres. (**e**) Histograms of diameter, deformation, and RI for the microsphere mixture with (red) and without (blue) numerical refocusing applied. Scale bars: 10 μm.

Further analysis from the refocused phase maps allowed quantitative characterization of the deformability, size, and RI of the microspheres. Because both PMMA and silica microspheres are rigid with the same diameter, the 2D scatter plot of deformation versus area displayed a similar distribution, which fails to distinguish between the microsphere population (Fig. 2c). However, the 2D scatter plot of RI versus diameter effectively differentiated the two populations (Fig. 2d). The measured RIs and diameters for PMMA and silica microspheres were *n* = 1.4934 ± 0.0034, d = 4.140 ± 0.125, and *n* = 1.4602 ± 0.0016, d = 4.149 ± 0.237 (mean ± SD), respectively (see Supplementary Table 1), in agreement with the manufacturer’s specifications. Histograms of deformation, diameter, and RI with and without numerical refocusing in Fig. 2e clearly show that the numerical refocusing significantly enhances the precision of quantitative characterization, as shown by narrower data distribution with reduced standard deviations of each measured quantity by 7% to 61% (see Supplementary Table 1 and Supplementary Figure 2). This substantial enhancement in measurement precision highlights the potential of QP-DC to contribute to more precise characterization of samples in imaging flow cytometry.

### Mass density characterization of blood cell types in whole blood

We further demonstrated the capability of QP-DC to characterize different blood cell types in whole blood from a healthy donor (Fig. 3). The acquired complex optical fields clearly reveal frequent blood cell types including erythrocytes, thrombocytes, lymphocytes, neutrophils and monocytes (Fig. 3a). The 2D scatter plot of deformation versus area effectively distinguished erythrocytes, thrombocytes, and lymphocytes (Fig. 3b). However, as neutrophils and monocytes have similar deformation and area, it is difficult to differentiate them, which is consistent with previous RT-DC studies^25^. In contrast, as shown in Figure 3c, the 2D scatter plot of RI versus area with the QP-DC measurements can easily distinguish each cell type based on their RI values. Since the RI value, *n*, of most biological samples is linearly proportional to mass density, *C*, as *n* = *n*_m_ + *αC*, where *n*_m_ is the RI of medium and *α* is the RI increment (*dn/dc*), the mass density of each population can be directly calculated. Here, we used *α* = 0.19 ml/g for most blood cells^26,27^ and *α* = 0.15 ml/g for erythrocytes^28^, reflecting the unique RI increment value of haemoglobin in erythrocytes. The measured mass density values are consistent with previous reports^29,30^, which validates our measurements (see Supplementary Table 2). The present implementation of QP-DC thus highlights its ability to quantitatively characterize both mechanical and biophysical parameters of biological samples, providing a robust method to differentiate different cell types among blood samples according to RI values.

**Figure 3.**
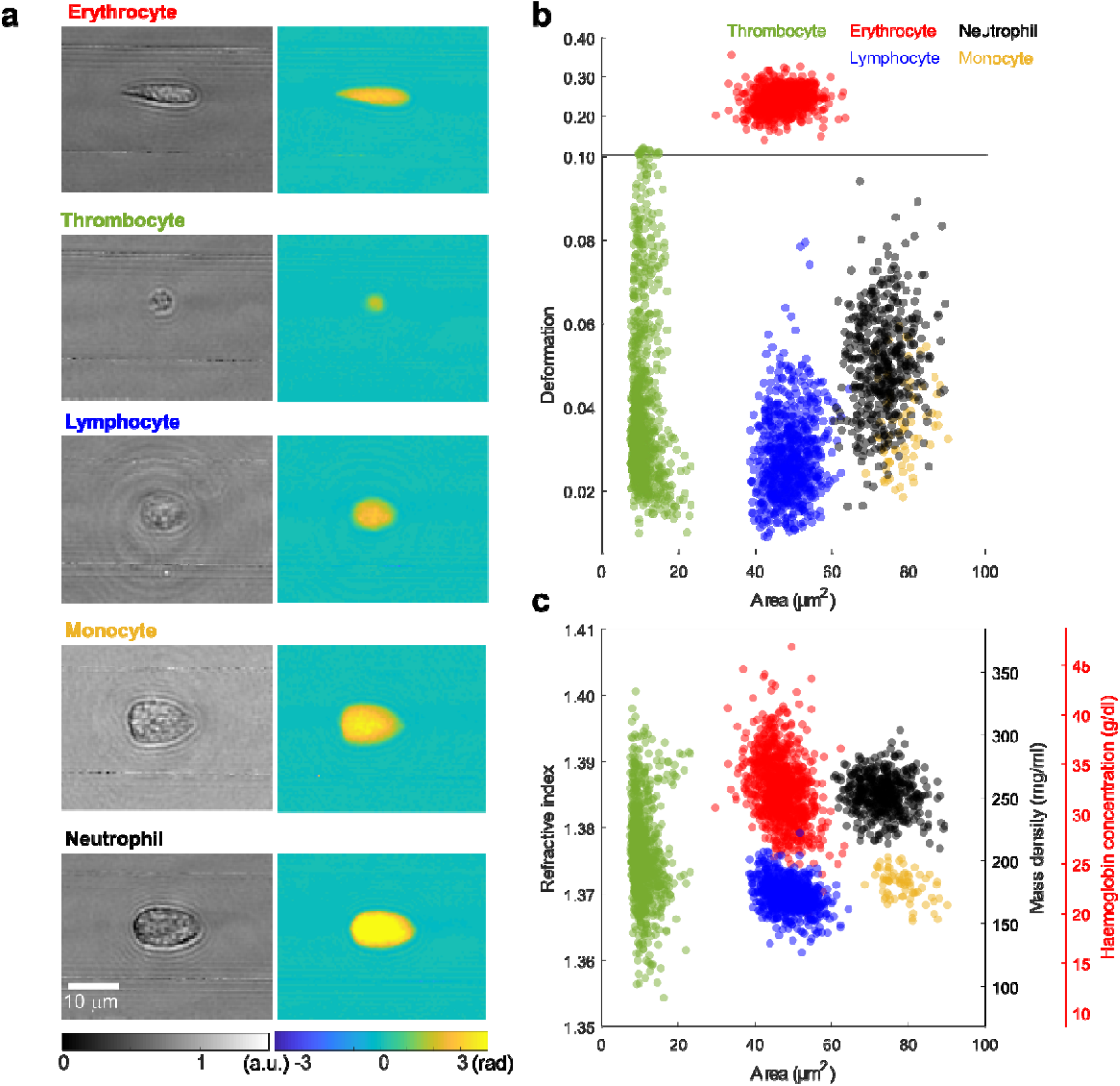
Quantitative phenotyping of blood. (**a**) Representative amplitude (left) and phase (right) maps of various blood cell types. (**b** – **c**) The 2D scatter plots of (**b**) deformation and (**c**) RI versus area. The color of each dot represents different blood cell types: thrombocytes (green), erythrocytes (red), lymphocytes (blue), neutrophils (black), and monocytes (gold). In (**c**), haemoglobin concentration for erythrocytes and mass density for all cell types are shown on a separate y-axes. The number of data points representing erythrocytes in (**b**) and (**c**) was intentionally reduced by a factor of 100 for clarity in visualization.

Conventional RT-DC studies have efficiently distinguished blood cell types using a 2D scatter plot of brightness versus cell area^25^. However, this approach requires precise adjustment of brightness to maintain the consistent contrast between samples and the background. More critically, conventional RT-DC requires intentional slight defocusing to extract cell contours from the thin halo around the cell, and this defocusing criterion may vary significantly among users^14^. In contrast, QP-DC utilizes differences in RI between blood cell types as an alternative feature for distinguishing each cell type. It is noteworthy that except for erythrocytes, where haemoglobin absorption predominates in the visible range^31^, other blood cells have similar absorbance^32^, and the 2D scatter plot of RI versus area closely resembles that of brightness versus area in conventional RT-DC measurements^25^. This observation is consistent with imaging models such as defocusing microscopy^33^ and the transport of intensity equation (TIE) imaging^34^, where intensity contrast is proportional to the focal offset and the Laplacian of phase delay. Consequently, in the underfocused condition^14^, samples with higher RI at similar cell diameter appear darker due to increased optical path length gradients. It suggests that the brightness distribution acquired in conventional RT-DC might result from RI difference at the defocused plane.

### Characterization of LPS-stimulated neutrophils

To further demonstrate the capability of QP-DC for biological and clinical studies, we investigated the mass density differences of neutrophil after activation. Isolated neutrophils from 6 healthy donors were activated with lipopolysaccharide (LPS) for 40 minutes to trigger a rapid response (see Materials and Methods). Previous studies have shown that LPS-mediated neutrophil activation results in significant morphological and rheological changes, including volume expansion and softening^12^. Our QP-DC measurements revealed that the deformation of LPS-stimulated neutrophils was 15.1 ± 9.2 % higher (mean ± SD) than that of unstimulated neutrophils, with a notably wider standard deviation in deformation distribution (Figs. 4a-d, j and Supplementary Table 3). The mass density of neutrophils from each healthy donor significantly decreased by an average of 3.6% following LPS stimulation. Specifically, the mass density dropped from 263.38 ± 9.27 mg/ml in unstimulated neutrophils to 254.08 ± 16.72 mg/ml in stimulated cells (Figs. 4e–h, k, and Supplementary Figure 3). This reduction was predominantly attributed to volume expansion while the dry mass remained unchanged (Fig. 4l).

**Figure 4.**
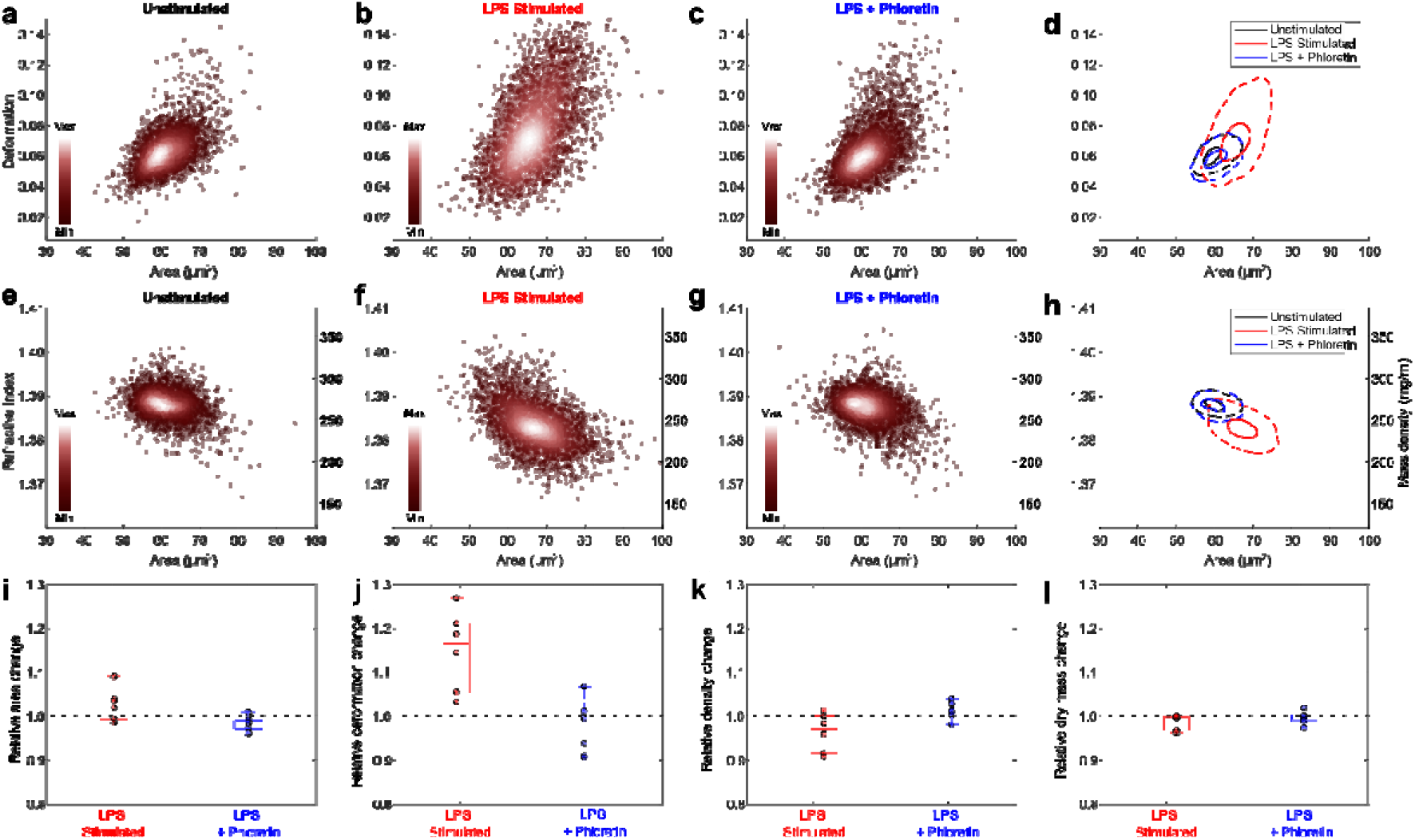
Quantitative characterization of neutrophils under lipopolysaccharide (LPS) stimulation. 2D scatter plot of deformation versus area for (**a**) unstimulated (**b**) LPS-stimulated, and (**c**) phloretin-preincubated neutrophils prior to LPS stimulation from a healthy donor. (**d**) Corresponding kernel density estimate plots illustrating the distribution differences in deformation and area for the three conditions shown in (**a – c**). 2D scatter plot of refractive index (RI) versus area for (**e**) unstimulated, (**f**) LPS-stimulated, and (**g**) phloretin-preincubated neutrophils prior to LPS stimulation. (**h**) Corresponding kernel density estimate plots illustrating the distribution differences in RI and area for the three conditions shown in (**e – g**). (**i** - **l**) Quantitative comparisons of (**i**) area, (**j**) deformation, (**k**) mass density, and (**l**) dry mass of LPS-stimulated, and phloretin-preincubated neutrophils prior to LPS stimulation. For each donor, parameters were normalized to the mean values of their own unstimulated neutrophils. Dashed lines indicate the mean values for the unstimulated condition. The number of healthy donors was 6.

Previous studies have shown that the volume expansion in LPS-stimulated neutrophils is mediated by water influx through aquaporins^12,35^. To investigate this further and study how neutrophil mass density is affected, we conducted control experiments with neutrophils incubated with phloretin, an inhibitor for aquaporin-9 to disrupt water influx^36^ before stimulation with LPS^37^. QP-DC measurements indicated that the mass density, dry mass, and deformation of neutrophils pre-incubated with phloretin were not affected by LPS stimulation (Figs. 4c, g, i-l). It confirms that inhibiting water influx effectively attenuates the activation of neutrophils initiated by LPS. These findings demonstrate that QP-DC can measure subtle mass density differences quantitatively in biological samples under physiological and pathological conditions.

### Characterization of low-density neutrophils in patients with SLE

The stimulated neutrophils with lower mass density, classified as LDNs, are increasingly recognized for their significant roles in immune response and immune-mediated disease, yet their origins and functions remain poorly understood^8,11^. SLE is an autoimmune disease with complex aetiology, in which LDNs are notably elevated and are believed to contribute to disease pathology and progression^9^. Conventionally, LDNs are detected when they float in the fraction of mononuclear cells after density-gradient centrifugation of whole blood. This is a time-consuming process and may alter neutrophil physiology. The diagnosis of SLE and its disease activity are commonly assessed using the SLE-DAI by trained clinicians^38^. However, the direct relationship between neutrophil biophysical properties and SLE-DAI scores is still elusive.

In contrast, QP-DC offers a rapid and quantitative alternative for assessing neutrophil biophysics in patients (Figure 5). Using QP-DC, we analysed whole blood samples from healthy donors and individuals with SLE without density-gradient centrifugation and preserves neutrophil native states, which is critical for clinical applications. Patients were grouped based on disease activity: those with an SLE-DAI score of 0 were categorized as “SLE inactive,” while scores above 0 were referred to as “SLE active.” QP-DC successfully revealed distinct differences in neutrophil deformation and mass density across the healthy, inactive, and active SLE cohorts (Figures 5a and 5c). In particular, the average mass density of neutrophils was significantly reduced in SLE patients: 245.46 ± 9.63 mg/ml (SLE inactive) and 228.75 ± 7.94 mg/ml (SLE active), compared to 261.28 ± 3.69 mg/ml in healthy controls. These changes were accompanied by volume expansion, as reflected in the increased projected area: 72.60 ± 1.04 μm^2^ in healthy donors versus 78.56 ± 6.03 μm^2^ and 83.26 ± 3.70 μm^2^ in the SLE inactive and active groups, respectively. These findings confirm the presence of LDNs in SLE (disease specific marker) and demonstrate a trend between decreased neutrophil mass density and disease severity (activity specific marker).

**Figure 5.**
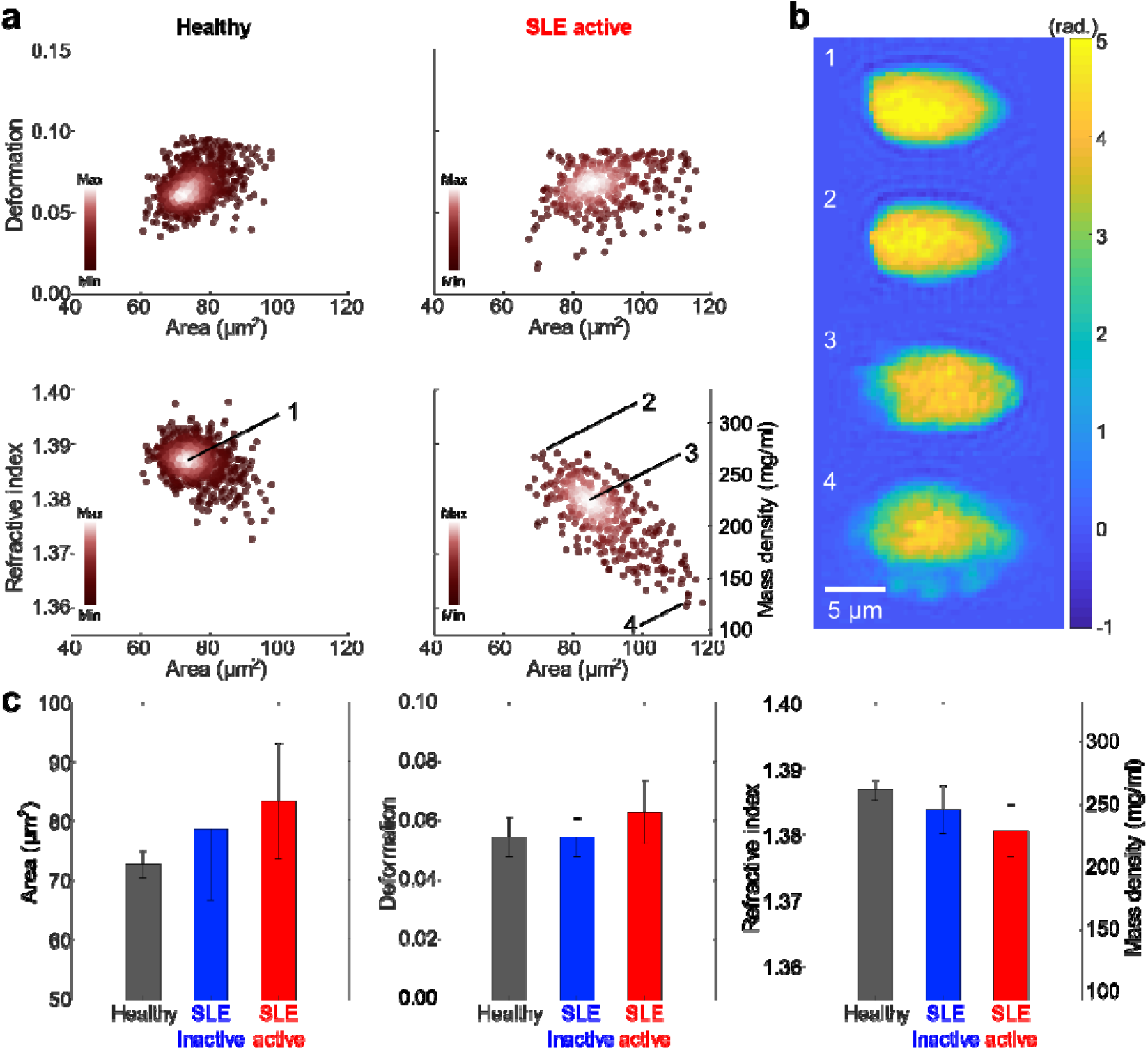
Quantitative characterization of low-density neutrophils (LDNs) in patients with systematic lupus erythematosus (SLE). (**a**) 2D scatter plots of deformation (top) and RI (bottom) versus area of neutrophils in whole blood from a healthy donor (left) and an patient with SLE (right). (**b**) Phase maps of representative neutrophils from the healthy donor (1) and the patient with SLE at various RI values (2 – 4). The corresponding data points for each neutrophil in the 2D scatter plot are indicated in (**a**). (**c**) Mean area, deformation, and RI of neutrophils from three populations: healthy donors (n = 5), patients with inactive SLE (SLE Disease Activity Index [SLE-DAI] = 0, n = 4), and patients with active SLE (SLE-DAI > 0, n = 7).

Representative phase images (Figure 5b) further support these observations. A neutrophil with higher mass density (cell “2”) in a SLE active patient appears morphologically similar to healthy controls (cell “1”), while those with reduced mass density (cells “3”and “4”) exhibited irregular surface features, indicating early stages of neutrophil extracellular trap (NET) formation^10,12^. Interestingly, although cell deformability increased only modestly (by 3.4%) in the SLE active group, this shift may still reflect disease-associated mechanical changes. The neutrophil in the SLE active group exhibiting the lowest mass density (cell 4 in Fig. 5b) displayed surface protrusions, which may represent either attached activated thrombocytes or apoptotic blebbing. During the image analysis, these protruding structures were included in the segmented cell contour, potentially leading to an overestimation of cell area and underestimation of mass density. Nonetheless, the central region of the cell showed reduced phase values and dry mass, which supports the reduced mass density in the SLE active group. Incorporating more advanced segmentation approaches could help distinguish protrusions from the main cell body, improving accuracy in such heterogeneous cases^39^. Overall, these results highlight the capability of QP-DC to characterize neutrophil heterogeneity in diseases like SLE. This suggests its potential utility in uncovering the mechanistic roles of LDNs in autoimmune pathophysiology, with implications for diagnosis and disease monitoring.

## Discussion

In this study, we developed and validated QP-DC, a novel technique that integrates deformability cytometry with QPI employing self-reference interferometry. This approach, validated with a mixture of microspheres and whole blood, demonstrates enhanced feature precision via numerical refocusing based on Rayleigh-Sommerfeld propagation, which corrects for axial position variations and ensures consistently focused images. Moreover, QP-DC not only measured conventional morphological parameters from imaging flow cytometry but also quantitatively characterized biophysical properties such as dry mass and mass density, which broadens the scope of cellular analysis.

Our study demonstrates the potential of QP-DC in characterizing changes in neutrophil mass density during LPS stimulation and in patients with SLE. We found that LPS-stimulated neutrophils have a lower mass density due to water influx, confirmed by inhibition experiments using phloretin. Additionally, we observed that the decrease in neutrophil mass density in LDNs correlates with disease severity in SLE. These results suggest that QP-DC can contribute valuable insights into neutrophil behaviour, potentially supporting clinical diagnostics and enhancing the understanding of immune-mediated diseases. The observed reductions in mass density and increases in cell area in SLE patients, particularly those with active disease, suggest that these biophysical markers correlate with disease severity and may reflect functional changes such as intravascular NET formation. Unlike traditional density-gradient methods, QP-DC preserves native cell states and offers a practical approach for monitoring disease-associated neutrophil phenotypes directly from whole blood. In future studies, larger cohort studies will be essential to validate the diagnostic and prognostic value of these parameters across different stages and subtypes of SLE and related chronic inflammatory rheumatic diseases. Integrating QP-DC with molecular profiling techniques may uncover mechanistic links between LDN biophysics and immune dysregulation, potentially guiding targeted therapeutic strategies.

The integration of QPI with deformability cytometry in QP-DC provides a distinctive approach compared to other recent developments of high-throughput QPI methods that merge QPI with microfluidics^40–43^. Holographic and ultrafast optofluidic implementations have enabled label-free classification of leukocyte subtypes and subcellular phase imaging of cultured cells at very high throughput, highlighting the versatility of QPI-based cytometry^44,45^. While these approaches provide valuable insights, QP-DC extends their scope by directly addressing immune cell mechanics in physiologically and clinically relevant contexts. By combining QPI with deformability cytometry, QP-DC enables simultaneous measurement of morphology, mechanics, and intrinsic biophysical parameters. Numerical refocusing from measured complex optical fields further ensures consistently focused images and more precise contour extraction, with future potential to incorporate machine learning for enhanced segmentation process^46^ and speed up the calculation of focused fields^47^. Beyond its technical strengths, QP-DC allowed us to detect reductions in neutrophil mass density under LPS stimulation and identify heterogeneous subpopulations in SLE, thus demonstrating its potential for translational applications in immune cell biology. We expect that the capabilities of QP-DC to provide additional biophysical parameters in a high-throughput manner can open future applications in drug screening and omics studies^48^, which can enrich our understanding of cell biomechanics. Moreover, we envision that with modified sample loading techniques, time-resolved cell mechanical analysis can be performed in a high-throughput manner.

To conclude, QP-DC combines QPI with deformability cytometry, enabling precise image-based analysis and assessment of biophysical features via numerical refocusing. This offers a precise and comprehensive characterization of cellular properties. We argue that QP-DC has the potential to become a widely used tool in both research and clinical diagnostics, and will contribute to our understanding of cell mechanics and diseases.

## Materials and Methods

### Sample Preparation

The 50 *μ*l of the mixture of 4-μm diameter poly(methyl methacrylate) (PMMA, 73371-5ML-F, Sigma-Aldrich) and silicon dioxide microsphere (54375-5ML-F, Sigma-Aldrich) are diluted in 950 *μ*l sucrose solution. In order to decrease the RI mismatch between microspheres and medium, 40% (w/w) sucrose/water solution is used, and the RI of the solution was measured by an Abbe refractometer (ORT1RS, Kern & Sohn GmbH).

### Isolation and activation of human neutrophils

With approval for the study (Ethikantrag Nr. 98_18 B; Nr. 334_18 B) from the ethics committee of the Uniklinikum Erlangen, we obtained blood from individual donors with their informed consent in accordance with the guidelines of good practice and the Declaration of Helsinki. Fresh whole blood was collected from patients with SLE or healthy human donors into EDTA tubes (S-Monovette® EDTA K3E, 9 ml, Sarstedt). To isolate neutrophils, we performed density gradient centrifugation at 350 g for 30 minutes at room temperature using Histopaque (Histopaque®-1077, Sigma-Aldrich). The high-density layer, resting above the red blood cell layer, underwent twice hypotonic erythrocyte lysis. The neutrophil pellet was then resuspended in Roswell Park Memorial Institute (RPMI) 1640 medium containing 2 mM Glutamine and 100 U/ml Penicillin-Streptomycin (all Gibco). The concentration of viable cells was determined using acridine orange/propidium iodide staining (Logos Biosystems) with the Luna-FL™ Dual Fluorescence Cell Counter (Logos Biosystems). Neutrophils were activated with 3 µg/ml LPS from Klebsiella pneumoniae (Sigma) for 20 min (or 40 min). For the inhibition assay, neutrophils were pre-incubated with 50 µM (or 10 µM) phloretin (Sigma Aldrich).

### Microfluidic chip

The microfluidic chip is made of polydimethylsiloxane (PDMS) fabricated in the TDSU Lab-on-a-chip systems in Max Planck Institute for the Science of Light, Erlangen, Germany. The microfluidic chip has a 300-μm-long narrow channel with a 20 μm × 20 μm square cross-section. The detailed information for the fabrication of microfluidic chips can be found elsewhere^4^. The sample suspension was loaded into a syringe attached to a syringe pump (Cavro Centris, Tecan Group Ltd.). The total flow rate was 0.06 *μ*L/s, of which the sheath flow rate was 0.045 *μ*L/s and the sample flow rate was 0.015 *μ*L/s. The total flow rate was monitored by a flow sensor (LG16-0430D, Sensirion AG) during the measurements.

### Quantitative phase deformability cytometry

The optical setup for QP-DC employs quantitative phase imaging based on self-reference interferometry^19,49^. Briefly, a collimated laser beam from a fibre-coupled laser diode (*λ* = 520 nm, LP520-SF15, Thorlabs Inc.) illuminates a sample flowing along the microfluidic chip. The scattered light from the sample is collected by an objective lens (40×, NA = 0.5, Carl Zeiss AG) and a tube lens (*f* = 150 mm). The sample image is further magnified by additional relay lens sets with the 4-*f* configuration and the total magnification is 90×. To measure complex optical field, self-reference interferometry consisting of a Rochon polarizer (MgF_2_, 1.5° beam separation, RPM10, Thorlabs) and a linear polarizer (LPVISE100-A, Thorlabs) is inserted along the optical path. The Rochon polarizer splits the beam into two, the ordinary and extraordinary beam with a separation angle, and two beams interfere at the image plane and generate a spatially modulated hologram. The hologram is recorded by a high-speed CMOS camera (CB013MG-LX-X8G3, XIMEA GmbH) with the frame rate of 3,000 Hz. In order to mitigate the motion blurring artefact for the samples flowing inside microfluidic channels with a high flow rate, the exposure time was reduced to 3 microseconds. All the experiments were conducted in the very narrow time window to exclude any influence of sample preparation and microscopy setup.

### Data analysis

The complex optical field was retrieved from the acquired spatially modulated hologram by applying a field retrieval algorithm based on Fourier transform^22^. For numerical refocusing, an axial stack of optical fields at various distances was compute numerically by employing the Rayleigh-Sommerfeld backpropagation. Among the axial stack, the optical field of a sample at the focal plane was selected according to a specified focus criterion.

In detail, the optical fields at arbitrary distances along the optic axis, denoted as E_z_(*r*), were calculated by applying the convolution of the retrieved field at the camera plane, E_0_(*r*) = *A*(*r*)exp(*iϕ*(*r*)) where *A*(*r*) and *ϕ*(*r*) are amplitude and phase map, respectively, with the Rayleigh-Sommerfeld propagator, 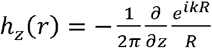, as E_z_(*r*) = E_0_ (*r*) * *h*_*z*_(*r*), where * is the convolution operator, *R*^2^=*r*^2^+*z*^2^ and *k*=2*πn*/*λ* is the wavenumber of light within a medium of refractive index *n*. This numerical propagation was efficiently conducted in the Fourier space as E_*z*_ (q) = ℱ{E_z_ (*r*)} = ℱ{E_0_ (*r*)} *H*_*z*_(*q*), where ℱ represents the Fourier transformation and 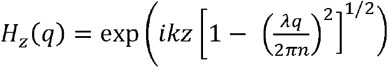is the Fourier transformation of the Rayleigh-Sommerfeld propagator^23,50^. The axial stack of propagated optical fields was generated, covering distances ranging from *z* = −10 μm to 10 μm with a step size of 0.2 μm. By calculating the gradient of amplitude maps at the boundaries of each sample for each axial plane, the axial plane exhibiting the lowest gradient value was identified as the focal plane.

In the focused optical field, the sample region was segmented from the background based on phase values. The deformation was calculated from the convex hull contour of the segmented region as 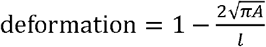, where *A* is the area enclosed by the convex hull contour and *l* is the length of the convex hull contour^4,51^. The volume of the cell, *V*, was estimated by rotating the contour with the symmetric axis under the assumption that the cells flowing in the microfluidic channel has the cylindrical symmetry. The mean RI of each sample,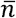, is calculated from the 2D summation of the focused phase map, *ϕ*(x,y), as 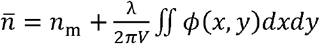, where is the RI value of surrounding medium. The dry mass of the sample, *m*, is calculated by 2D summation of the focused phase map inside the segmented region, as 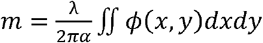, where *α* is the refractive index increment (dn/dc) with *α* = 0.15 ml/g for erythrocytes^28^ and *α* = 0.19 ml/g for other blood cells^26,27^.

## Acknowledgements

We thank S. Girardo, S. Khan, S. García Rey, and L. Strampe (MPL) for assisting microfluidic control and fruitful discussion. We thank P. Patel, part of the TDSU Lab-on-a-chip systems at MPL, for the production of the PDMS microfluidic chips. We acknowledge financial support from the AutoRAPID (Project number: 40-01366-2170-0001 to K.K., E.O., F.M., P.R., P.L., M.K., J.L., and J.G.), National Research Foundation of Korea (NRF) funded by the Ministry of Science and ICT (No.2022K1A3A1A04062969 to J. Sh.) and Max Planck Society core funding (to J.G.).

## Author contribution

K.K. and J.G. conceived the project. K.K., C.S., J.Sc., and J.Sh. performed experiments. C.S., J.Sc., J.R., and M.H. prepared biological samples. F.M., P.R., P.L., and J.L. developed microfluidic control. K.K. and E.O. analysed data. M.K. and M.H. participated in critical discussions. K.K. wrote the manuscript. All authors reviewed the manuscript.

